# The integration of continuous audio and visual speech in a cocktail-party environment depends on attention

**DOI:** 10.1101/2021.02.10.430634

**Authors:** Farhin Ahmed, Aaron R. Nidiffer, Aisling E. O’Sullivan, Nathaniel J. Zuk, Edmund C. Lalor

## Abstract

In noisy, complex environments, our ability to understand audio speech benefits greatly from seeing the speaker’s face. This is attributed to the brain’s ability to integrate audio and visual information, a process known as multisensory integration. In addition, selective attention to speech in complex environments plays an enormous role in what we understand, the so-called cocktail-party phenomenon. But how attention and multisensory integration interact remains incompletely understood. While considerable progress has been made on this issue using simple, and often illusory (e.g., McGurk) stimuli, relatively little is known about how attention and multisensory integration interact in the case of natural, continuous speech. Here, we addressed this issue by analyzing EEG data recorded from subjects who undertook a multisensory cocktail-party attention task using natural speech. To assess multisensory integration, we modeled the EEG responses to the speech in two ways. The first assumed that audiovisual speech processing is simply a linear combination of audio speech processing and visual speech processing (i.e., an A+V model), while the second allows for the possibility of audiovisual interactions (i.e., an AV model). Applying these models to the data revealed that EEG responses to attended audiovisual speech were better explained by an AV model than an A+V model, providing evidence for multisensory integration. In contrast, unattended audiovisual speech responses were best captured using an A+V model, suggesting that multisensory integration is suppressed for unattended speech. Follow up analyses revealed some limited evidence for early multisensory integration of unattended AV speech, with no integration occurring at later levels of processing. We take these findings as evidence that the integration of natural audio and visual speech occurs at multiple levels of processing in the brain, each of which can be differentially affected by attention.

## Introduction

In everyday life, face-to-face speech communication is a multisensory experience. Seeing a speaker’s face – especially under challenging listening conditions – can greatly improve speech comprehension (Sumby and Pollack 1954; Grant and Seitz 2000; Ross et al. 2007). A process known as multisensory integration – wherein information from different senses is integrated by the brain to form a coherent representation of the speech (Stein and Stanford 2008) – is thought to drive this improvement in comprehension (Campbell 2008). Another factor strongly influencing speech comprehension is our ability to selectively attend to one of many competing sources of speech – known as the cocktail-party phenomenon (Cherry 1953). In real-world environments, the targets of attention and multisensory integration are often aligned (i.e., when we want to understand what our conversational partner is saying). However, sometimes they are not (i.e., when we are eavesdropping). How attention and multisensory integration interact in these contrasting scenarios is not well understood.

In general, much debate surrounds the role of attention in multisensory integration. According to one account, audiovisual integration can occur automatically and independent of attention. For instance, audiovisual integration underlying the ventriloquism effect (the illusion of a sound coming from the location of a concurrent visual cue other than the sound’s true location) has been demonstrated to be unaffected by selective attention (Bertelson et al. 2000; Vroomen et al., 2001). Indeed, in such cases the interaction can often be better thought of in reverse, with pre-attentively integrated audiovisual stimuli recruiting attention in a bottom-up manner (Driver 1996; Olivers and Van der Burg 2008). However, this notion of bottom-up attentional capture by automatically integrated multisensory stimuli has been challenged by a number of other studies reporting modulatory effects of top-down attention on multisensory integration. For instance, audiovisual integration was shown to be reduced when attention was diverted to a secondary task (Alsius et al. 2005; 2007). Besides, EEG research has suggested that selective attention can affect event-related brain potentials (ERP) related to multisensory processing and the effects are larger for attended compared to unattended stimuli (Talsma and Woldorff 2005; Talsma et al., 2007). Attempts to reconcile these varied findings have been put forth. According to one perspective (Talsma et al. 2010), how much the objects/events in the environment compete with each other in terms of salience may determine the nature of the interaction between attention and multisensory integration; less competitive, salient stimuli could be integrated automatically and pre-attentively whereas competitive stimuli require top-down attention to be integrated. This multifaceted interplay may be due to the fact that audiovisual integration takes place at multiple levels of processing in the brain (Eskelund et al., 2011; Peelle and Sommers 2015; Baart et al. 2014), and the interaction between attention and multisensory integration may depend on the level of processing at which the integration occurs (Koelewijn et al., 2010).

More work is needed to test this idea of differing interactions across different processing levels. As mentioned above, this is particularly true for the case of natural, continuous audiovisual speech, which has been the focus of relatively little multisensory integration research (but see Fairhall and Macaluso 2009; Morís Fernández et al. 2015). Indeed, a majority of the studies informing us about multisensory integration and how it interacts with attention have either involved non-speech stimuli such as beeps and flashes (Senkowski et al. 2005; Fujisaki et al. 2006) or simple speech stimuli such as syllables, isolated words or short segments of speech (Alsius et al. 2014; Tiippana et al., 2004). And, where the work has centered on speech, it has mostly focused on the well-known McGurk effect (McGurk and MacDonald 1976) — an illusory speech sound resulting from exposure to mismatched auditory and visual syllables. However, everyday speech communication consists of a dynamic flow of meaningful, connected words rather than discrete syllables or isolated words, along with a variety of facial and articulatory gestures that accompany the ongoing acoustic information. Therefore, although the use of syllabic stimuli has been very enlightening, it is not sufficient to allow us to understand how cocktail-party attention and multisensory integration interact in the context of the kind of audiovisual speech we encounter in our daily lives.

An opportunity for progress on this issue comes from recent progress in the development of methods for studying the neural processing of natural, continuous speech, and how that processing is affected by attention (Ding and Simon, 2012; Mesgarani and Chang, 2012) and multisensory input (Zion Golumbic et al. 2013; Luo et al. 2010). This includes work from our group showing strong effects of both attention (Power et al. 2012; J. A. O’Sullivan et al. 2015) and visual speech (Crosse et al., 2015; 2016a) on the cortical tracking of various dynamic speech features. But studies investigating how attention and multisensory integration interact in this context have been lacking. This is the goal of the present study. Specifically, we aim to apply a framework for indexing the audiovisual integration of natural speech (Crosse et al., 2015; 2016a) to data acquired during a naturalistic multisensory cocktail-party experiment (O’Sullivan et al. 2019). In doing so, we find stark differences in electrophysiological indices of audiovisual speech integration for attended vs unattended audiovisual speech.

## Results

The main focus of our study was the influence of attention on natural, continuous audiovisual speech integration. To explore this question, our primary dataset was EEG recorded during a multisensory cocktail-party experiment (O’Sullivan et al. 2019). This experiment involved two conditions: 1) where subjects attended to an audiovisual speaker while ignoring a second, spatially separated audio-only speaker (attended condition), and 2) where subjects attended to the audio-only speaker while continuing to look at the audiovisual speaker but ignoring what he was saying (unattended condition).

Our primary analysis of the resulting data involved modeling responses to the audiovisual speech when it was attended versus when it was ignored. However, for reasons that will be explained below, our modeling of this primary dataset was carried out based on models fit using a second data set. That second data set was collected – using the same EEG system and 128-channel montage – while a different set of subjects was presented with speech (in noise) from a single speaker in 1) an audio-only condition, 2) a visual-only condition (i.e., silent lipreading), or 3) an audiovisual condition. Critically, the models derived from this second dataset were applied in precisely the same way to both the attended and unattended conditions in our primary cocktail-party dataset, allowing us to conduct an unbiased test of the influence of attention on audiovisual speech integration.

### Modeling audiovisual speech integration in a cocktail-party experiment

Following a long line of multisensory research (Stein and Meredith 1993; Besle et al., 2004), our strategy for modeling audiovisual speech integration was based on testing the hypothesis that audiovisual speech processing (AV) is not simply the linear sum of audio and visual speech processing (A+V). In the context of natural continuous speech, this can be done by modeling EEG responses to audiovisual speech and comparing the performance of such AV models to summed models that were fit separately to audio speech and visual speech (A+V). We have previously done this for single speakers in quiet (Crosse et al., 2015) and in noise (Crosse et al., 2016a). Importantly, this approach requires having EEG responses to audiovisual, audio-only, and visual-only stimuli – something that we did not have for our multisensory cocktail-party experiment. As such, we derived our models from one of these previously published datasets – specifically the one involving a single-speaker in noise (Crosse et al., 2016a).

This speech-in-noise dataset was acquired from twenty-one subjects who listened to a single speaker in spectrally matched noise presented in audiovisual (AV), audio-only (A) and video-only (V) conditions. To quantify how the EEG relates to the speech, we used linear regression to fit backward models (known as decoders) that aim to reconstruct the speech envelope from the EEG response it produced (Crosse et al. 2016b). We did this separately for the audio, visual and audiovisual speech conditions to get separate A, V and AV decoders for each subject (as in Crosse et al., 2015, 2016a). The unisensory A and V decoders were then summed to form A+V decoders for each subject (Fig 1a; see Materials and Methods for further details). The AV and A+V decoders were then averaged across all subjects to produce one generic AV decoder and one generic A+V decoder. Channel weightings for these generic AV and A+V decoders show the importance of contributions from both temporal and occipital scalp in relating the EEG data to the speech envelope (Fig. 1A right). Armed with these decoders, multisensory integration in the cocktail party can then be assessed by comparing how well the speech envelope can be reconstructed from audiovisual EEG using an AV decoder vs a summed A+V decoder. Any improvement in performance using the AV decoder can be attributed to multisensory processing. This is how we analyzed our primary dataset – from the multisensory cocktail-party experiment (Fig 1B).

**Figure 1.**
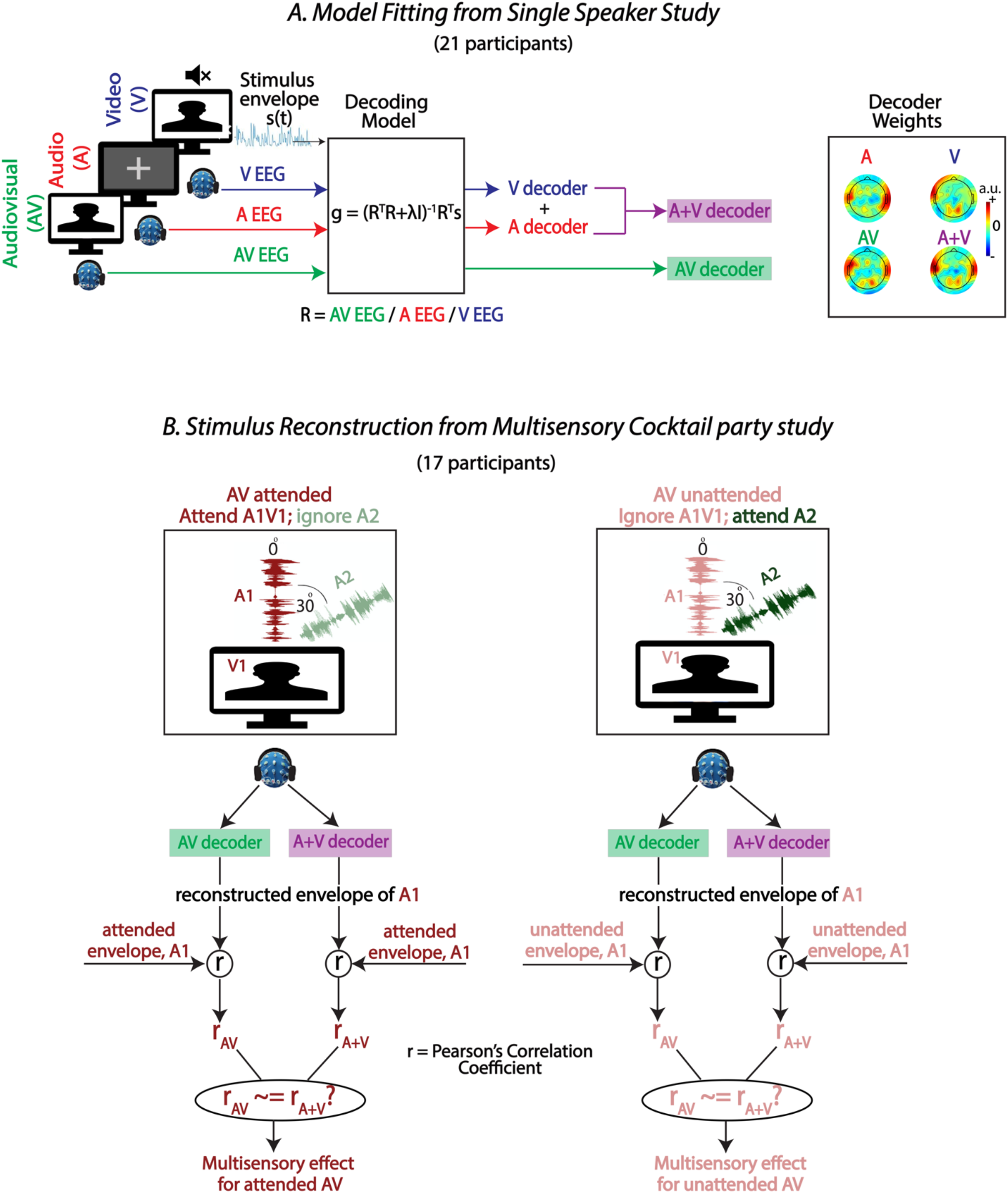
Experimental procedure and analysis approach. Silhouette images are used to protect identity of the speakers. **A**. Subjects were presented with a male speaker in −9dB noise in audiovisual (AV), audio-only (A) and video-only (V) format. Envelope reconstruction models AV and (A+V) decoders were derived from their EEG recordings. The channel weightings for each decoder averaged across time-lags from 0 to 500 ms and across all subjects (N = 21) are shown on the right. **B**. A separate set of subjects performed a multisensory cocktail-party task where they were presented with an audiovisual speaker (A1V1) directly in front of them and an audio only speaker (A1) at 30° to their right (over headphones). They had to either attend or ignore A1V1. The AV and A+V decoders obtained from the single-speaker paradigm were applied to reconstruct the envelope of the audiovisual speaker (A1) when he was attended vs unattended. Multisensory integration in both conditions was quantified as the difference between reconstruction accuracy (Pearson’s r between the actual and reconstructed envelope of A1) using the AV decoder (r_AV_) and A+V decoder (r_A+V_).

The multisensory cocktail-party experiment involved seventeen subjects (different from the participants in the speech in noise dataset) who listened to two speakers where one of the speakers was presented audio-visually (A1V1) while the other one was presented, with 30° of spatial separation, in the auditory modality only (A2). In one condition, subjects had to pay attention to the audiovisual speaker while ignoring the audio-only speaker. In the other condition, they had to pay attention to the audio-only speaker while ignoring the audiovisual speaker. Importantly, in both conditions, subjects were asked to maintain fixation on the face (specifically the eyes) of the audiovisual speaker displayed on a screen (Fig. 1B; see Materials and Methods for details).

To assess compliance in the attention task, subjects were required to answer between four and six multiple-choice questions (each with 4 possible answers) on *both* stories after every 1-minute trial. As previously reported for this dataset (O’Sullivan et al. 2019), subjects successfully performed the task, answering 76.1 ± 9.7% of questions correctly on the A1V1 content when it was attended (significantly greater than 25% chance, one-sided Wilcoxon signed-rank test, *p* = 1.6 × 10^−4^). This dropped to only 30.3 ± 9.1%, when A1V1 was not attended, although this was still significantly greater than 25% chance (*p* = 0.0277), reflecting the difficulty of completely ignoring an AV speaker that one is looking at. This difficulty was also reflected in the performance of subjects in attending to the A2 speaker, which was 61.4 ± 12.8% (significantly greater than 25% chance, *p* = 1.6 × 10^−4^), but lower than the A1V1 attended performance (*p* = 2.9 × 10^−4^). On the contrary, ignoring the A2 speaker when attending to A1V1 was relatively easy, reflected in subjects not performing better than chance in answering questions on the A2 content when A1V1 was attended (26.4 ± 4.8%, *p* = 0.14). We also used an eye tracker to monitor subject compliance with our request to fixate on the speaker’s eyes. Again, as previously reported (O’Sullivan et al. 2019), subjects looked at the eyes (as instructed) more than the mouth of the speaker (A1V1 attended – percent time viewing eyes: 30.4 ± 15%, percent time viewing mouth: 11.9 ± 11.2%; *p* = 0.02; A1V1 unattended – percent time viewing eyes, 40.9 ± 23%, percent time viewing mouth: 5.8 ± 10.2%; *p* = 6.1 × 10^−4^). There was a slight difference between conditions in these viewing patterns, however, with subjects tending to look more at the mouth during the A1V1 attended condition compared with A1V1 unattended condition (Wilcoxon signed-rank test, *p* = 0.035).

The behavioral results above clearly show that attention had a significant effect on which speech stream subjects understood. To assess how this attention effect influences multisensory integration, we used our generic, pre-fit AV and A+V decoders to reconstruct the speech envelope of the audiovisual speaker from the EEG when the A1V1 speaker was attended and when he was unattended. Crucially, while the generic decoders were fit on data from other subjects, they were both applied in the same way during each condition – always focused on decoding the audiovisual speaker’s speech, with the only difference being whether the decoded data were acquired when that speaker was attended or ignored. Again, any difference in the reconstruction accuracy arising from the AV and A+V decoders points to a non-additive multisensory process.

### EEG responses to attended, but not unattended, audiovisual speech are better modeled as a multisensory process

We first verified that our generic decoders could accurately reconstruct the envelope of the audiovisual speaker when he was attended and ignored. This was assessed by calculating the Pearson’s correlation between the reconstructed envelope and the actual envelope of the audiovisual speaker in both conditions, and chance was established using permutation testing. At the group level, the reconstruction accuracies were significantly better than chance for both AV and A+V decoders in both attention conditions (AV decoder: attended condition, *p* = 1.60×10^−4^, unattended condition, *p* = 6.43×10^−4^; A+V decoder: attended condition, *p* = 1.92×10^−4^, unattended condition, *p* = 3.88×10^−4^; one-sided Wilcoxon signed-rank test against chance-level; Fig. 2A). These results suggest that, overall, the neural tracking of speech envelope is robust across individuals. Therefore, although the decoders were trained on a different set of subjects performing a different task, they were able to decode the speech envelope from the multisensory cocktail-party EEG. Moreover, the reconstruction accuracies were significantly greater for the attended condition than the unattended condition for both AV and A+V decoders (AV decoder: *p* = 6×10^−4^; A+V decoder: *p* =0.003; two-sided Wilcoxon signed-rank tests), in line with the stronger neural representation of attended speech in EEG (O’Sullivan et al. 2015).

**Figure 2.**
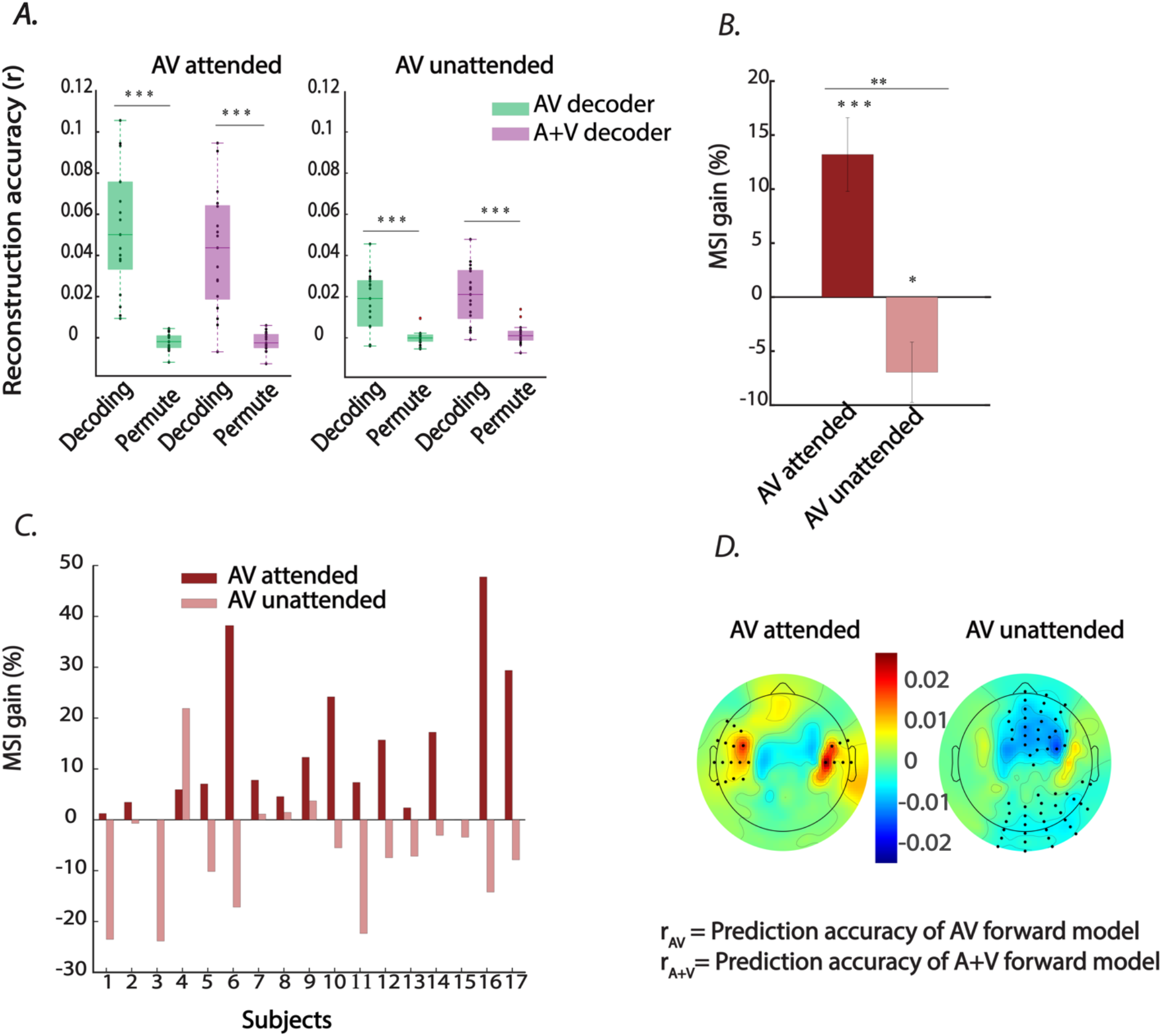
Robust multisensory integration when AV speech is attended vs unattended. **A**. Both attended (left) and unattended (right) audiovisual speech envelope could be reconstructed significantly better than chance (determined by permutation test). **B**. Grand-average (N = 17) normalized multisensory gain was positive for attended, and negative for unattended audiovisual speech. **C**. Normalized multisensory gain at single-subject level. The sharp dissociation across attention (more positive multisensory gain for attended, than for unattended; trial-averaged) was visible for all but subject 4. **D**. Topographical distribution of EEG prediction accuracies (obtained from forward modeling). The black markers indicate channels where the multisensory integration effect [AV – (A+V)] was significant across subjects (p < 0.05, two-sided cluster-based permutation test). *** p < 0.001, ** p < 0.01, * p < 0.05.

But 1) do the reconstruction accuracies from the AV and A+V decoders differ from each other? 2) does the measure of multisensory integration vary across attention conditions? The answer to both of these questions was yes. We found that the reconstruction accuracy based on the AV decoder was significantly higher than the A+V decoder (i.e., AV-[A+V] > 0) for the attended condition (*p* = 7.13×10^−4^; one-sided Wilcoxon signed-rank test), which suggests that responses to attended audiovisual speech are not simply the sum of independent audio and visual speech processing, i.e., they include nonlinear multisensory contributions. In contrast, we found that the reconstruction accuracy based on the AV decoder was significantly lower than the (A+V) decoder (i.e., AV-[A+V] < 0) for the unattended condition (*p* = 0.0168; one-sided Wilcoxon signed-rank test). In other words, the data in the unattended condition are better modeled under the assumption that they derive from two separate, noninteracting, unisensory (A+V) processes. This suggests a relative lack of multisensory integration for unattended audiovisual speech. Indeed, the very fact that the AV decoder performed worse for unattended speech suggests that any multisensory processes contributing to the generic decoder were actively harmful in modeling the unattended audiovisual speech. To directly compare the magnitude of multisensory integration across all subjects, we derived a summary measure of “multisensory gain” for each subject based on the normalized difference in performance between the AV and A+V decoders (i.e., (AV-(A+V)) / (|AV| + |A+V|)). This was done to ensure that the index of multisensory integration was not biased by the variability in correlation values across conditions as well as across subjects. Across subjects, there was a sharp dissociation in multisensory gain across attention conditions. In particular, the multisensory gain for attended audiovisual speech was significantly greater than zero (*p* = 2.30×10^−4^, one-sided Wilcoxon signed-rank test), while multisensory gain for unattended audiovisual speech was significantly less than zero (*p* = 0.0079, one-sided Wilcoxon signed-rank test). Accordingly, the multisensory gain for attended audiovisual speech was significantly greater than that for unattended audiovisual speech (*p* = 0.001; two-sided Wilcoxon signed-rank test; Fig. 2B). Furthermore, this sharp dissociation of multisensory gain (more positive for attended than unattended) was visible for all but subject 4(Fig. 2C).

How might we gain insight into the potential neural regions driving the effects of attention on our decoder-based multisensory gain metric? It can be difficult to interpret the weights of multivariate decoders (Haufe et al. 2014). However, one can also build so-called forward encoding models that seek to capture how the EEG responses reflect various stimulus features, and these can lend themselves to clearer interpretation. To that end, we used regularized linear regression on the data from the single speaker study to produce forward encoding models (also known as temporal response functions, Crosse et al. 2016b) that reflect how the EEG on each channel was affected by the speech envelope. We did this for the audiovisual condition to produce an AV forward model. And we did it for the audio-only and visual-only conditions to produce A and V forward models, respectively, that we then summed to produce an A+V forward model. We then used these AV and A+V forward models to predict EEG responses from the cocktail party dataset. Topographical distributions of the prediction accuracies (Pearson’s r) averaged across all subjects, are shown in Figure 2D. We found that, in the attended condition, the AV model outperformed the A+V model over focal regions of left and right temporal scalp (Fig. 2D, left), suggesting multisensory integration in bilateral temporal brain regions. In the unattended condition, the A+V model outperformed the AV model (or, perhaps more intuitively, the AV model underperformed the A+V model) across fronto-central and parieto-occipital scalp. We found these topographical distributions to be fairly consistent across majority of the subjects (Supplementary figure 1).

### Temporal analyses of model performance suggest different interactions of attention and multisensory integration at different levels of processing

Previous work has suggested that the interaction between attention and multisensory integration may depend on the level of processing at which the integration occurs (Koelewijn et al., 2010). Our encoding and decoding analyses, meanwhile, seem to show a fairly simple dissociation with strong evidence for multisensory integration to attended speech and an apparent lack of integration for unattended speech. However, these analyses were carried out by relating the speech envelope to the EEG using a broad range of stimulus-EEG time-lags (0 – 500 ms, see Materials and Methods). This misses an opportunity to examine the effects of attention on multisensory integration at different time-lags – as a proxy measure of processing at different cortical stages.

To explore this idea, we first trained decoders aimed at reconstructing the speech envelope from the single-speaker EEG data at individual time-lags between 0 and 500 ms. We then used these single-lag decoders to reconstruct the speech envelope using the cocktail-party dataset. As before, we first tested if each of the single-lag decoders could reconstruct speech envelope significantly better than chance. We found that the AV decoder was able to reconstruct both attended and unattended speech envelope and the A+V decoder was able to reconstruct the attended speech envelope significantly better than chance at all of the single time-lags (all *p* < 0.01; permutation tests). However, the A+V decoder was not able to reconstruct the unattended speech envelope from time-lags between 0 – ∼100 ms (all *p* > 0.05; permutation tests).

In the attended condition, the AV decoder significantly outperformed the A+V decoder at a broad range of time-lags from 0 – ∼180 ms, and again at time-lags from ∼420 – 500 ms (Figure 3A). And the A+V decoder never outperformed the AV decoder. This fits with our earlier results in showing that attended audiovisual speech responses are best modeled when including multisensory processing contributions. However, the story for the unattended condition is more nuanced. Consistent with our results above, the A+V decoder generally outperformed the AV decoder at time-lags longer than ∼170 ms, indicating that, at these longer latencies, EEG responses to unattended audiovisual speech are better modeled as the sum of separate audio and visual unisensory processes. However, in contrast to the earlier decoding results, at shorter time-lags (0 – 100 ms), the AV decoder actually outperformed the A+V decoder, providing evidence of some multisensory integration for unattended audiovisual speech at early stages of processing. Moreover, the finding that the A+V decoder did not perform better than chance at these early time-lags supports the notion that some multisensory integration is happening in this window, which our unisensory additive A+V model was not able to capture. This early effect (AV > A+V) was also present in the attended condition, although it was significantly bigger than that observed in the unattended condition (*p* = 0.0016; two-sided Wilcoxon signed-rank test), suggesting that the early multisensory integration effect is further strengthened by attention.

**Figure 3.**
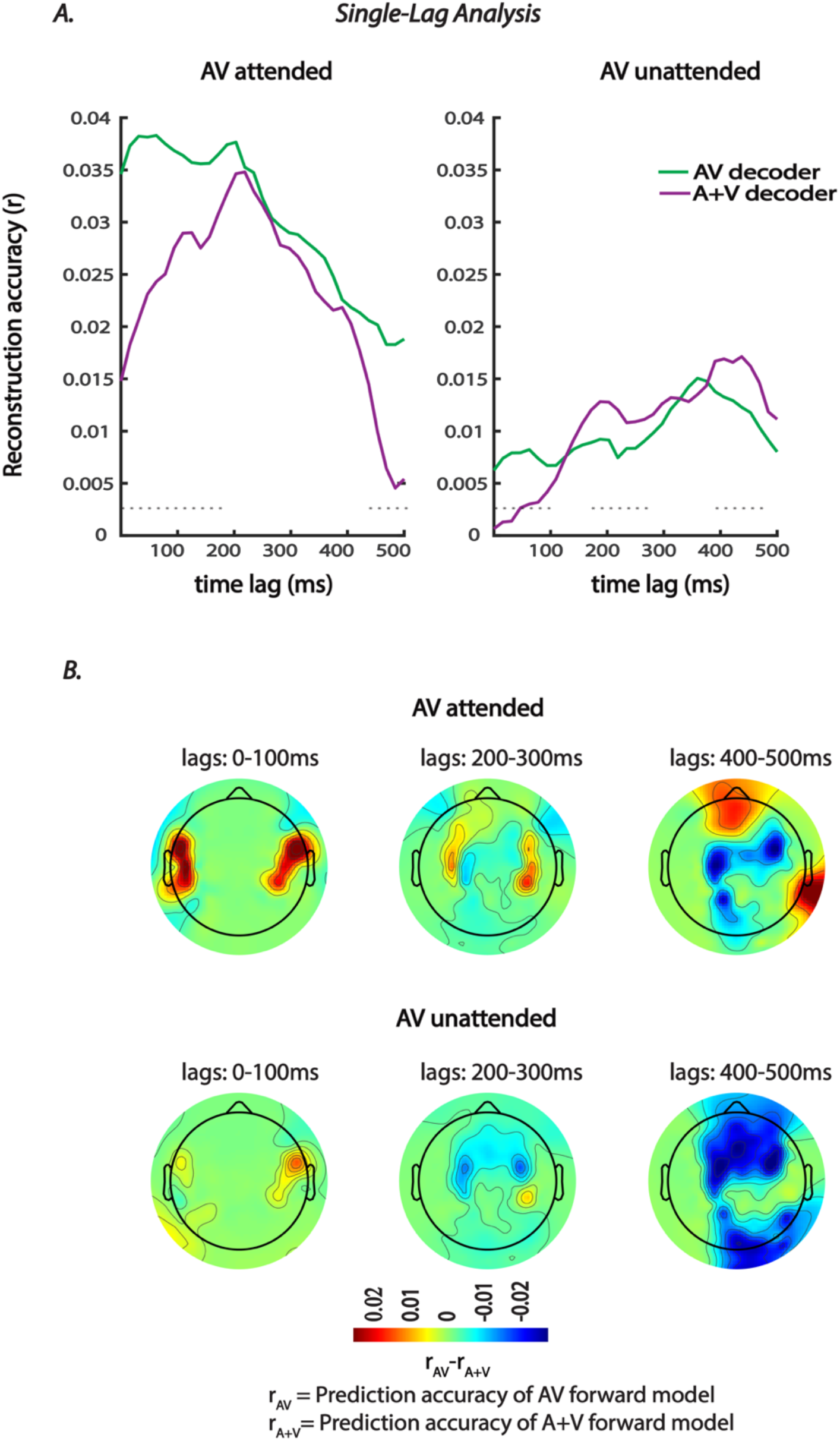
Spatiotemporal analysis reveals more complex dissociation of multisensory interaction across attention conditions. **A**. Single-lag reconstruction accuracies obtained using AV and (A+V) decoders at every lag between 0 and 500 msec. The black markers running along the bottom of each plot indicate time lags when the multisensory integration effect [AV-(A+V)] was significant across subjects (p<0.05, two-sided cluster-based permutation test). **B**. Topographic distributions of EEG prediction accuracies at specific time lags.

This interpretation was supported by the topographic distribution of EEG predictions using single time-lag forward modeling (Figure 3B). The previously seen focal bilateral advantage for the AV model was visible for attended speech from 0 – 300 ms. And this same pattern was visible (albeit more weakly) for the unattended speech from 0 – 100 ms. Thereafter, the unattended speech over front-central and parieto-occipital scalp was best modeled as a sum of the two unisensory processes. This suggests the possibility of multisensory integration for both attended and unattended audiovisual speech at early processing stages (before 100 ms), with continued integration occurring at higher levels only for the attended speech.

### No clear relationship between the performance of models based on speech envelopes and behavior

In our multisensory cocktail party experiment, our two attention conditions attempted to simulate two aspects of real-world behavior, i.e., looking at one’s conversational partner while attending to them versus continuing to look at them (so as not to cause offense) while eavesdropping on another conversation. As reported above, performance on the multiple-choice questions when attending to the audiovisual speaker was consistently high (76.1 ± 9.7%). Attending to the audio-only speaker was more difficult, which was reflected in lower scores and more variability across subjects (61.4 ± 12.8%). We hypothesized that, the better job the subjects do at attending to the audiovisual speaker (as reflected in their performance on the questions), the greater their multisensory integration. Therefore, we expected to see a positive correlation between multisensory integration and behavioral performance for attended audiovisual speech. However, we failed to find a significant correlation (Pearson’s r =0.28, *p* = 0.28). On the other hand, when attending to the audio-only condition, we hypothesized that the better the subjects are at answering questions on the attended story, the better the job they are doing in suppressing the audiovisual speaker and the lower should be their multisensory integration. Therefore, we expected their multisensory integration to be negatively correlated with their behavioral performance in the unattended audiovisual condition. Again, we failed to find a significant correlation (r = −0.34, *p* = 0.12).

## Discussion

We have demonstrated that when attending to audiovisual speech, cortical responses to the speech envelope are better captured by a multisensory model (AV) than the sum of two independent unisensory processes (A+V), reflecting integrative processes. This was true across a broad range of time-lags and was especially pronounced over bilateral temporal scalp regions. In contrast, when attention is directed away from an audiovisual speaker, EEG responses to that speaker are, overall, best modeled as the sum of separate unisensory audio and visual processes. Spatiotemporal analysis of this result revealed some evidence for multisensory integration over bilateral temporal scalp regions at short latencies (< 100 ms), with no clear evidence for integration at later time lags. Together, these results provide evidence that AV speech integration occurs at multiple stages, each of which can be differentially affected by attention.

Our findings appear to agree with previously published perspectives arguing that multisensory integration can happen in parallel at different levels of processing (Calvert and Thesen 2004; Schwartz et al., 2004; Eskelund et al., 2011; Peelle and Sommers 2015), and that attention can influence integration at different levels depending on the task and the nature of the stimuli (Talsma et al. 2010; Koelewijn et al., 2010). When the amount of competition between the stimuli is low (for example, if one stimulus is more salient than the other), multisensory integration tends to occur automatically, whereas top-down attention is required for multisensory integration when multiple stimuli compete for processing resources. Here, to test the interaction between multisensory integration and attention, we have used a cocktail-party paradigm which is an ideal example of competing stimuli. The audiovisual speaker and the concurrent audio-only speaker were both competing for neural processing and therefore, according to Talsma (2010), top-down attention is required for multisensory integration. If multisensory integration occurred regardless of attention, the face and the voice of the audiovisual speaker would still be integrated throughout the unattended condition, and we would have observed similar multisensory gain across both attention conditions. The fact that we observed such a stark difference in our multisensory integration index across the two conditions (Fig. 2B, C) indicates that multisensory integration is very sensitive to attentional focus in the context of natural speech.

We sought to explore the idea that attention might influence multisensory integration differently at different hierarchical levels of processing. To do this, we used single-time analysis at different latencies as a proxy measure of processing at different hierarchical levels. In doing so, we found that the interaction with attention appears to vary at different levels. Although the A+V model outperforms the AV model for the unattended audiovisual speech when combining neural data across a broad range of time lags (0-500 ms), the opposite appears to be the case at early time lags between 0 – 100 ms. This suggests some audiovisual enhancement at shorter latencies, even though subjects were not paying attention to the audiovisual speaker. We interpret this finding based on a wealth of evidence that has amassed to show that multisensory integration can happen automatically and without the need for attention (i.e., pre-attentively) based on shared low-level spatial or temporal properties (Van der Burg et al. 2011; Tang et al., 2016; Atilgan et al. 2018). The temporal dynamics of auditory and visual speech are quite strongly correlated (Chandrasekaran et al. 2009). Thus, this correlation may have driven some early, attention-independent integration of the auditory and visual speech for the unattended audiovisual speaker in a bottom-up manner. This early effect was also visible in the attended condition although it was much larger than the unattended condition. We conclude that congruent audio and visual events lead to some multisensory integration at early latencies regardless of attention, but when attention is also directed to the multisensory stimulus, this early integration effect is further enhanced. This is consistent with the notion of multiple stages of interaction between multisensory integration and attention (Talsma and Woldorff 2005), and with the general idea that selective attention improves the representation of low-level stimulus features (Maddox et al. 2015). Beyond 100 ms, there is no clear evidence of multisensory integration for the unattended speech – at least according to our approach based on comparing multisensory and unisensory models. These findings are in line with proposals that early sensory areas may automatically integrate signals to increase their bottom-up salience, but that attention may play a larger role in higher order association areas where signals are integrated into multisensory representations (Macaluso and Driver 2005; MacAluso et al. 2016)

This notion that attention exerts a stronger influence at later levels of processing is consistent with a long history of research on cocktail-party attention. From early behavioral work (Treisman, 1964), to more recent noninvasive (Power et al. 2012; Teoh & Lalor, 2020) and invasive (J. O’Sullivan et al. 2019) electrophysiology research, there is a substantial amount of support for the notion that attention may exert weaker effects at lower levels of processing, before leading to a marked separation in the strength of representation for attended and unattended speech at higher levels. Indeed, some of the EEG/MEG work that has been done on this question has specifically identified strong cocktail party attention effects at temporal latencies beyond 100 ms with more limited effects at shorter latencies (Power et al. 2012; Ding and Simon 2012; Puvvada and Simon 2017). If cocktail-party attention exerts weaker effects at low levels of processing, this explains why our data appear to show evidence of multisensory integration at short latencies for both attended and unattended speech. However, beyond 100 ms, we only see evidence of multisensory integration for attended speech, as the representation of the unattended speech “object” has likely been suppressed.

Because of the limited spatial resolution of EEG, it is difficult to draw definitive conclusions about the involvement of different anatomical regions in our results. However, we can attempt some inferences based on linking our findings to previous research using technologies with better spatial resolution. For example, both electrocorticography (ECoG) and functional magnetic resonance (fMRI) have implicated superior temporal regions (STG and STS) in the processing of speech beyond simply its low-level acoustics (Mesgarani et al. 2014; Hickok 2000). ECoG research over the last few years has also highlighted STG, in particular, as being an area where attended, but not unattended speech, is strongly represented (Mesgarani and Chang 2012; J. O’Sullivan et al. 2019). Meanwhile, both ECoG and fMRI have highlighted the STS as playing a critical role in integrating higher level features of audio and visual speech (Beauchamp et al. 2004; Zhu and Beauchamp 2017). In the present study, the superiority of the AV model over the A+V model in predicting attended AV data was most pronounced over a limited set of EEG channels focally located bilaterally over temporal scalp. This would seem to be consistent with attention-multisensory integration interactions deriving from superior temporal cortex. A notably different spatial pattern was apparent for the unattended speech, especially at longer latencies. In particular, the A+V model outperformed the AV model across fronto-central and parieto-occipital scalp regions. This difference in performance may have come about because using a multisensory decoder to model what are likely two separate unisensory (visual and auditory) processes, led to suboptimal predictions over scalp regions sensitive to those unisensory processes (parieto-occipital scalp, visual speech; fronto-central scalp, auditory speech). Future work using ECoG in a paradigm like ours could potentially confirm these suppositions.

Both multisensory integration (Sumby and Pollack 1954; Ross et al. 2007) and cocktail-party attention (Cherry 1953) strongly influence speech comprehension. Indeed, our behavioral results showed clear effects of attention on the ability of subjects to answer comprehension questions on the two speech streams. While attending to the audiovisual speech appears to have been easier, there was still a degree of variability in performance across subjects. This variability was even more pronounced in the more difficult eavesdropping condition, with lower overall performance on the questions, and even above-chance performance on the audiovisual speaker content when it was unattended. Indeed, it is likely that some (if not all) subjects will have found it difficult to maintain attention completely on a single speech stream at all times, particularly when attending to the audio-only speech and trying to ignore the very salient audiovisual speaker. It is possible that attention will have drifted between speech streams more for some subjects than others, contributing to the behavioral variability we see. Given the marked interaction between attention and multisensory gain in our EEG data, we expected to see some correlation between EEG-based multisensory gain and behavior on the attention task across subjects. Although the data appeared to trend in the predicted directions, neither of the two correlations we examined were significant. There are several potential explanations for this. First, cortical fold configuration and skull thickness will vary across subjects, such that different cortical generators will be represented with different levels of strength on each person’s scalp. We attempted to normalize our multisensory gain measure to account for individual differences in response magnitude, but this normalization will not necessarily have accounted for the different relative contributions of different generators. Second, our decoders were trained on a different set of subjects not performing the multisensory cocktail-party task. Although the decoders were able to reconstruct speech significantly better than chance, they may not be sensitive enough to capture nuanced neural activity patterns related to individual differences in behavioral performance on the cocktail-party task. Third, our behavioral measure of attention is imperfect. Asking questions on the speech content after every 1-minute trial is demanding on working memory as well as attention. And working memory abilities will have varied across subjects. Furthermore, some subjects may have been better than others at intuiting certain answers despite not having attended to the speech content. A final issue with respect to the behavioral variation across subjects and the fact that, overall, performance on the audiovisual speech content was greater than chance when that speaker was (supposedly) unattended. One possibility is that bottom-up capture by the audiovisual speaker may have caused some unwanted attentional drifting to that speaker. Indeed, this may be the reason we see early integration effects for the unattended condition. However, these early effects are at latencies that are much shorter than those in which we typically see strong cocktail-party attention effects (Power et al. 2012). So, linking them directly to high-level task behavior seems tenuous. Furthermore, the early attention-multisensory integration effect we see seems to be quite consistent across subjects (supplementary fig 2), even in the face of large behavioral variability across those same subjects. Future work utilizing more instantaneous measures of cocktail party attention would be helpful in adding further insight to this issue.

There are a number of other ways the work in the present study could be developed in future. As already mentioned, ECoG recordings could shed more light on the spatiotemporal details of the attention and multisensory interactions we have seen. Again, ECoG has been used for both attention (E. M. Zion Golumbic et al. 2013) and audiovisual speech (Ozker et al. 2018) research. Combining the two could be very powerful. In terms of EEG/MEG work, as well as using latency as a proxy measure for the “level” of speech processing, one could also explicitly explore how different acoustic, visual, and linguistic representations of audio and visual speech might be influenced by attention (Teoh and Lalor 2020) and multisensory interactions (O’Sullivan et al. 2020). To be able to confidently disentangle EEG responses indexing different representations would likely require optimizing the signal-to-noise ratio of the multisensory modeling framework we have used here. This would benefit from fitting individualized audio, visual and audiovisual decoders for each subject that takes part in the multisensory cocktail party study – something that we did not have in the present study. That said, the generic decoders used in the present study were eminently capable of modeling the data from a different experiment (Fig 2A) and revealed clear and interpretable differences in multisensory integration across attention conditions (Fig 2, 3). In the interests of highlighting that this result was not a foregone conclusion – and in the interests of full disclosure – we would also like to note here that we also attempted to model the multisensory cocktail-party data using generic decoders that were fit to a different AV dataset, specifically one that involved speech-in-quiet (the one used in Crosse et al., 2015). That analysis did not produce any coherent set of results across the attention conditions in the multisensory cocktail-party data. We take this as further supporting the interpretations we have drawn from the results presented above. This is because multisensory integration effects are known to be substantially larger for speech-in-noise than speech-in-quiet according to the principle of inverse effectiveness (Crosse et al., 2016a). Therefore, training the models on a dataset that is best able to capture the multisensory benefit arising from audiovisual speech gives them the power to reveal the multisensory integration-attention interactions in a multisensory cocktail-party. On the contrary, training the models on any dataset that does not fully leverage the benefit of multisensory integration will not produce the results we have seen here. Finally, extending the work to include even more naturalistic settings with multiple competing speakers could facilitate the testing of ideas around how competition and salience play into attention-multisensory interactions.

## Materials and Methods

### Subjects

Twenty-one subjects (6 females; age range 21-35 years) participated in the single speaker-in-noise experiment. This study was approved by the Ethics Committee of the Health Sciences Faculty at Trinity College Dublin. Seventeen different subjects (8 males; age range 18-30 years) took part in the multisensory cocktail-party experiment. This study was approved by the Research Subjects Review Board at the University of Rochester. Both studies were undertaken in accordance with the Declaration of Helsinki. All subjects were native English speakers, had self-reported normal hearing and normal or corrected-to-normal vision, had no history of neurological diseases and provided written informed consent.

### Stimuli and procedure

The stimuli used in the single speaker-in-noise experiment consisted of a male speaking on contemporary social, political and economic issues. Fifteen one-minute-long speech passages were rendered into 1280 x 720-pixel videos at 30 frames per second. The soundtracks were sampled at 48 kHz with 16-bit resolution, matched in intensity by root mean square normalization and mixed with spectrally matched stationary noise with SNR of −9 dB. Each passage was presented 3 times in audio-visual (AV), audio-only (A) and visual-only (V) format. The condition from trial-to-trial was randomized to prevent each passage from repeating in another format within 15 trials of the preceding one. Recording took place in a dark, sound-attenuated room and subjects were instructed to fixate on the speaker’s mouth during AV and V conditions, and a crosshair during A condition. Subjects had to detect target words during each trial via button press and rate the intelligibility of the speech stimuli at the end of each trial (the results of the target word detection task and speech intelligibility scores were not analyzed in this study).

In the cocktail-party experiment, subjects were presented with two competing male speakers (both different from the speaker in the previous experiment) narrating two classic works of fiction (story 1: Journey to the Centre of the Earth and story 2: Twenty Thousand Leagues under the Sea, both by Jules Verne). One speaker was presented in the audiovisual modality at 0° with no head related transfer function applied to the audio (A1) and the accompanying videos of the speaker were displayed on the monitor (V1) directly in front of the participant. The audio of the other speaker was convolved with a head related transfer function (taken from the CIPIC Database, Algazi et al. 2001) to be presented only in the auditory modality at 30° to the right (A2). There were two conditions depending on whether the audiovisual speaker was attended or not: 1) AV attended— subjects were instructed to pay attention to the audiovisual speaker (A1V1) while ignoring the concurrent audio-only speaker (A2). 2) AV unattended— subjects were instructed to attend to the audio-only speaker (A2) while continuing looking at the face the other speaker (V1) but ignoring the corresponding audio (A1). It is to be noted that to attend the audio-only speaker, subjects must ignore the competing audiovisual speaker and therefore, the condition is named AV unattended. Subjects undertook twenty 1-min long trials per condition and answered four to six multiple-choice questions on both stories after each trial.

### EEG acquisition and preprocessing

In both experiments, EEG data were acquired at a 512 Hz sampling rate from 128 scalp electrodes and two mastoid electrodes using an ActiveTwo system (BioSemi). Subsequent preprocessing was performed offline in MATLAB. The data were bandpass filtered between 0.3 and 30 Hz, downsampled to 64 Hz and referenced to the average of the mastoid channels. To detect channels with excessive noise, the time series from the single-speaker experiment were visually inspected in Cartool (brainmapping.unige.ch/cartool) and the standard deviation of each channel was compared with that of the surrounding channels. For the cocktail-party data, EEGLAB channel rejection methods such as spectrogram, kurtosis and probability distribution (Delorme and Makeig 2004) were used. In both experiments, noisy channels were recalculated by spline interpolation in EEGLAB.

To relate the speech stimulus to the EEG, the acoustic envelopes of the speech signals from both experiments were extracted (except for A2 in the cocktail-party experiment which we did not analyze here). The stimuli were first bandpass filtered into 128 logarithmically spaced frequency bands between 100 and 6,500 Hz using a gammatone filter bank. Then the envelope at each of the 128 frequency bands was extracted using a Hilbert transform, and the broadband envelope was obtained by averaging over the 128 narrowband envelopes. Afterwards, the envelopes and the EEG data were z-scored.

### Data analysis

Our goal was to investigate how multisensory integration is affected by attention. To do so, we quantified multisensory integration (MSI) using the additive model criterion (Stein and Meredith 1993) and assessed how that measure varied across attentional states in a multisensory cocktail-party environment. According to this criterion, multisensory integration can be indexed by the differences between cortical responses to multisensory stimuli (AV) and the summation of unisensory responses (A+V). To accomplish this, we used linear decoding models to reconstruct an estimate of the audio speech envelope from multichannel EEG (Crosse et al. 2016b). We trained the linear models (decoders) separately for audio, visual, and audiovisual speech conditions from the EEG data of the single speaker in noise experiment to produce separate A, V and AV decoding models. The algebraic sum of the A and V decoders was calculated to obtain an A+V decoder, representing independent unisensory processing. We then tested the performance of the AV and (A+V) decoders on the multisensory cocktail-party data in reconstructing the audio speech envelope of the audiovisual speaker from both attended and unattended conditions. The difference between envelope reconstruction accuracies of the AV and (A+V) decoders quantified multisensory integration in both conditions. It is important to note that no training was done on the cocktail-party data. The linear model employed to reconstruct the speech envelope, *s(t)* from the neural data, *r(t)* can be expressed as follows (Crosse et al. 2016b):

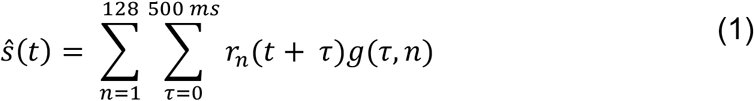

where *ŝ(t)* is the reconstructed envelope, *r*_*n*_ *(t* + *τ)* is the EEG response at channel n and time lag *τ* and *g*(*τ,n)* is the linear decoder for the corresponding channel and time lag. The time lags *τ* ranged from 0 to 500 msec poststimulus. The decoders from the single speaker-in-noise paradigm were obtained using ridge regression as follows:

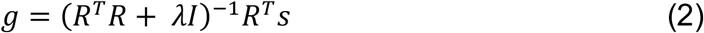

where *R* is the lagged time series of the EEG data, *λ* is the ridge parameter, *I* is the identity matrix and *s* is the speech envelope. If the EEG used to train the decoder *g(τ, n)* is from the AV condition of the single speaker in noise paradigm, then the resulting decoder is an AV decoder. Accordingly, if the EEG comes from the A-only and V-only conditions, then the resulting decoders are A decoders and V decoders respectively.

To get the AV decoder, leave-one-trial-out cross validation was used to choose the *λ* value (over a range of 2^4^, 2^5^,…, 2^14)^ that maximized the correlation between *ŝ(t)* & *s(t)* without overfitting to the training data. To get the A+V decoder, the algebraic sum of the A and V decoders was calculated for every combination of *λ* values (*λ*_*A*_ *=* 2^4^, *2*^5^,…, *2*^14^*; λ*_*V*_ *=* 2^4^, 2^5^,…, 2^14^). Each additive model was then used to reconstruct the envelope from the EEG data of corresponding AV condition to ensure that the model could reconstruct envelope from channels encoding both auditory and visual information. Leave-one-trial-out cross validation was again used to choose the combination of *λ* values that maximized the correlation between actual and reconstructed envelope. The process was repeated for all 21 subjects resulting in 21 AV decoders and 21 A+V decoders. Each decoder is the best envelope reconstruction model of a particular subject’s neural data. These decoders were then averaged across subjects to obtain a single AV decoder and a single A+V decoder which were then applied on the multisensory cocktail-party data

### Quantifying multisensory integration

The envelope (A1) of the audiovisual speaker (A1V1) in the cocktail party paradigm was reconstructed using equation (1) where *r* was the EEG data from either AV attended or AV unattended condition, *g* is either the AV decoder or the (A+V) decoder. Multisensory integration was then quantified for both conditions as:

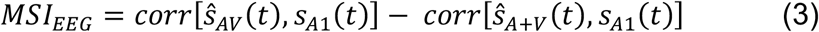

To compare the MSIs across the subjects, we normalized the difference metric so that the index is bounded between −1 and 1.

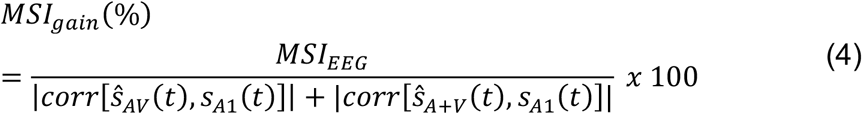

### Single-lag analysis

Previously, the AV and A+V decoders were integrating EEG data across a broad range of time lags from 0 to 500 ms. This was to make sure that important temporal information in the EEG can be captured to relate to each sample of the stimulus. However, there might be specific contribution from each time-lag towards attention-multisensory interaction that could be reflected in the envelope reconstruction accuracies. To investigate this, we trained AV and A+V decoders at individual lags from 0 to 500 ms, rather than integrating across them. As before, the decoders were averaged across subjects to produce one generic AV decoder as well as one generic A+V decoder at each time lag. Using the range of time lags from 0 to 500 ms at a sampling frequency of 64 Hz generated 33 individual lags and 33 separate AV decoders as well as 33 separate A+V decoders. Each of these decoders were then used to reconstruct the attended speech envelope (A1) of the multisensory cocktail-party data.

### Forward model

Forward models (also known as encoders) linearly transform the speech envelope to the recorded neural response at each channel location. While decoder weights are not neurophysiologically interpretable, forward model parameters are, with nonzero weights appearing at channels where cortical activity relates to stimulus encoding (Haufe et al. 2014). This allows us to investigate scalp topography to identify scalp regions that pick up the attention-multisensory interaction effects. To that end, we trained AV and A+V forward models at time lags between 0 – 500 ms using regularized linear regression (Crosse et al. 2016b). As done with the decoders, we also averaged the forward models across subjects to obtain a generic AV forward model and a generic A+V forward model which were then used to predict the EEG response to the audiovisual speaker in the cocktail-party data.

### Statistical Analysis

To test the performance of AV and A+V decoders on the cocktail party data against chance, nonparametric permutation tests were conducted (Combrisson and Jerbi 2015). A null distribution of 1,000 Pearson’s *r* values was created for each subject by finding the correlation between randomly permuted trials of predicted audio envelopes and actual audio envelopes. This was done for each decoder and for each attention condition separately. Permutation tests were also conducted in the similar way to test if the single-lag decoders were able to reconstruct speech envelope significantly better than chance. The threshold for above chance performance was *p* = 0.05 for each test. The true mean correlation coefficient served as the observed value of the test statistic. Comparison across conditions (attended vs unattended) and across decoders (AV vs A+V model performances) was conducted using Wilcoxon signed-rank test. Multiple comparisons were corrected for using nonparametric cluster-based permutation test (Maris and Oostenveld 2007)

## Acknowledgements

The work was supported by NIH grant DC016297 to ECL. Additional support was provided by a Science Foundation Ireland Career Development Award (CDA/15/3316) to ECL.

