## Supplementary Material for "The integration of continuous audio and visual speech in a cocktail-party environment depends on attention"

### AV attended

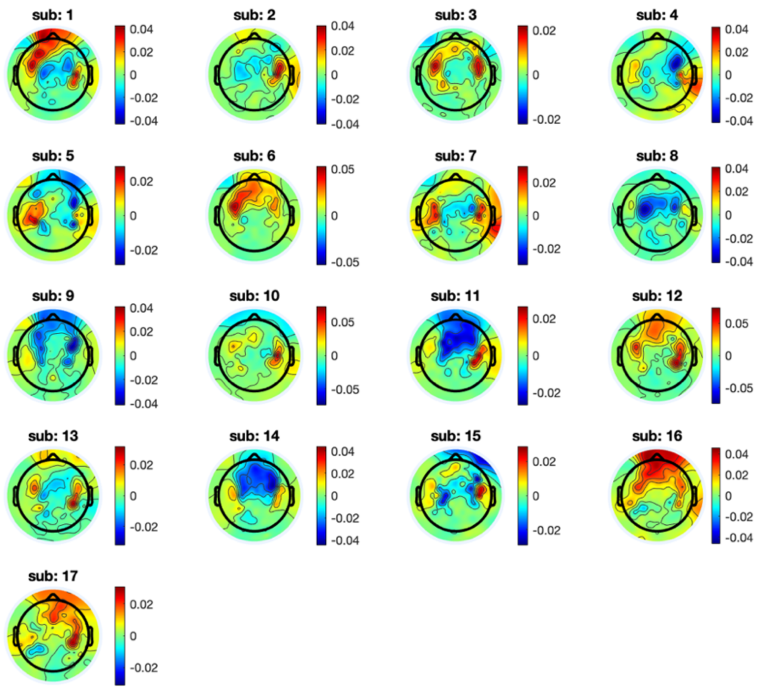


### AV unattended

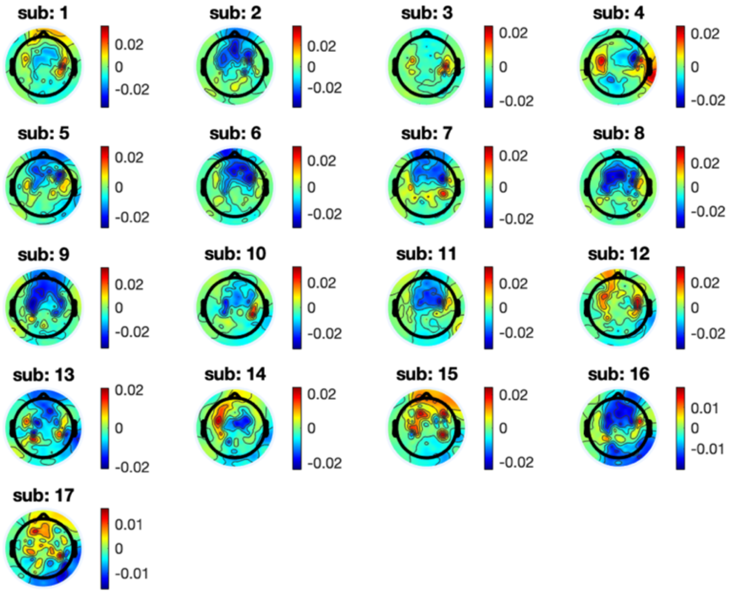


**Supplementary figure 1 | Topographical distribution of the EEG prediction accuracies, AV-A+V across 0-500 ms time-lags (related to Figure 2D).** For AV attended, multisensory benefit (AV> A+V) over bilateral temporal regions is clearly visible for subjects 1,3,4,5,6,7,12,13,14,15. For AV unattended, reduction of multisensory processing (AV < A+V) over fronto-central and parieto-occipital scalp) is clearly visible for subjects 1,2,5,6,7,8,9,10,11,13,14,16.

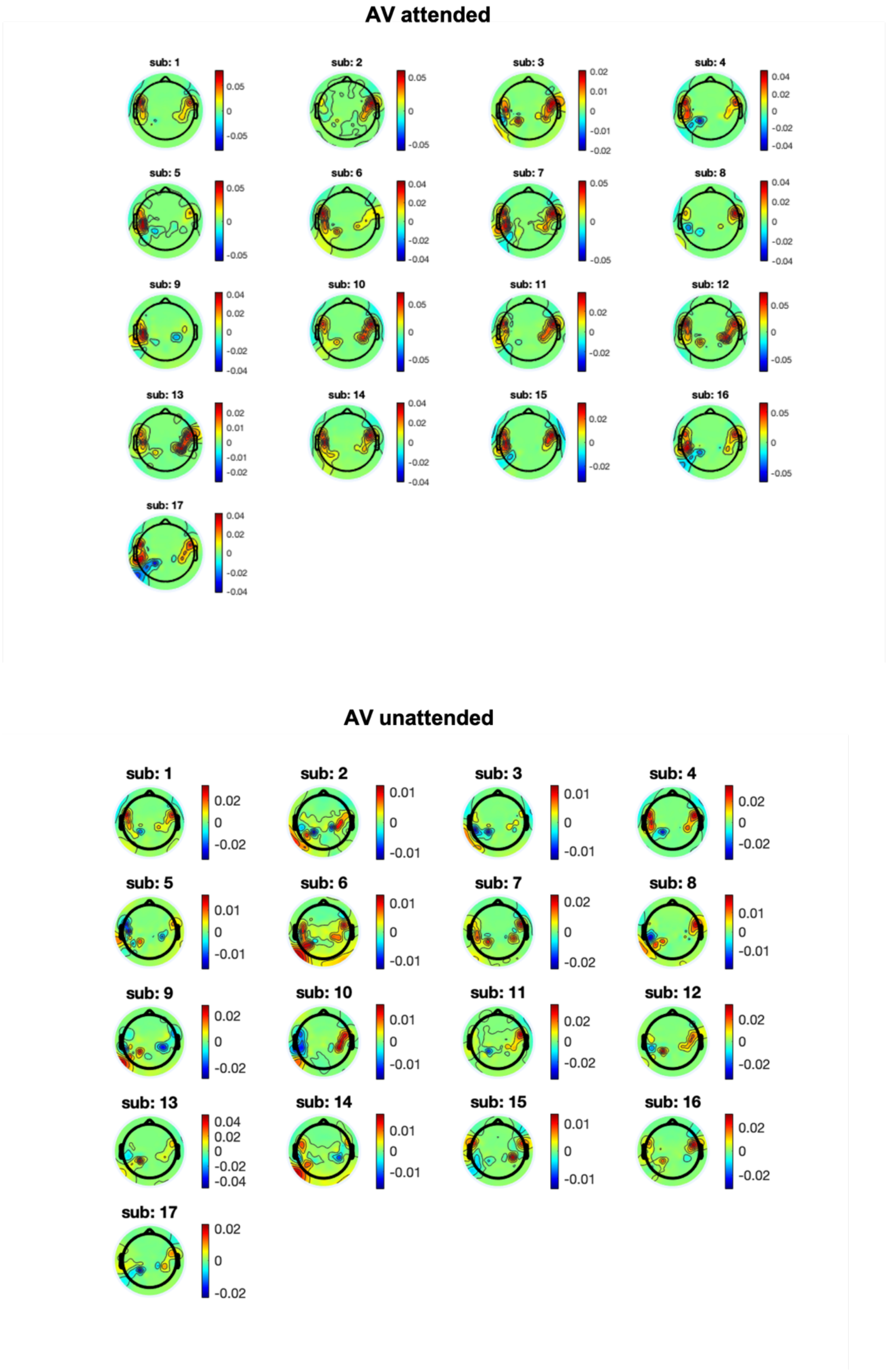


**Supplementary figure 2 | Topographical distribution of the EEG prediction accuracies, AV-A+V across 0-100 ms time-lags (related to Figure 3B).** Multisensory benefit (AV> A+V) at these time-lags over bilateral temporal regions is clearly visible for almost all subjects, suggesting early multisensory integration taking place for both attended and unattended audiovisual speech.
